# Towards understanding the bacterial biosynthesis of branched GDGTs: Identification of iso-diabolic acid-based tetraester and mixed ether/ester, membrane-spanning lipid intermediates in members of the Bacillota

**DOI:** 10.64898/2026.02.09.704826

**Authors:** Diana X. Sahonero-Canavesi, Nicole J. Bale, Melissa Antony Venancius, Michel Koenen, Ellen C. Hopmans, Jaap S. Sinninghe Damsté, Laura Villanueva

## Abstract

Branched glycerol dialkyl glycerol tetraethers (brGDGTs) are bacterial membrane-spanning lipids (MSLs) resembling archaeal membrane lipids, as they form monolayers and are linked to glycerol backbones via ether bonds. Ubiquitous in soils, sediments, and aquatic environments, their distributions are widely applied as paleoclimate proxies for reconstructing past temperature and pH. Despite this, understanding of their biological origins and functional role in cells remain incomplete. While some Acidobacteria are known producers of brGDGTs, genomic evidence and environmental surveys indicate additional bacterial contributors. In this study, we report the first detection of potential brGDGT biosynthetic intermediates in Bacillota. Ultra-high pressure liquid chromatography high resolution multi-stage mass spectrometry (UHPLC-HRMS^n^) revealed membrane-spanning diglycerol lipids which contained *iso*-diabolic acid (13,16-dimethyl octacosanedioic acid)-derived alkyl chains. These diglycerol lipids displayed diverse structures, including tetraesters, mixed ester/ether combinations, and vinyl ether bonds. Additionally, ‘open’ membrane spanning lipids analogous to brGTGTs were also identified. Notably, all brGDGT and brGTGT analogues were detected with a phosphatidylglycerol head group. Experiments showed that the two Bacillota strains, which produce these brGDGT biosynthetic intermediates, responded differently to changes in temperature and oxygen availability, suggesting that environmental regulation of brGDGT-related lipids is taxonomically dependent. Based on these findings, we propose a biosynthetic pathway for brGDGT formation and highlight the physiological implications for interpreting brGDGT-based paleoclimate proxies. This work expands the known diversity of bacterial sources of brGDGTs and provides new insights into the ecological and evolutionary significance of these lipids.

**IMPORTANCE:** Branched GDGTs (brGDGTs) are bacterial membrane spanning lipids which form a monolayer, linked through ether bonds to the glycerol backbone, characteristics more commonly found in archaeal membrane lipids. They are commonly used in paleoclimate proxies to assess past temperature and pH but their predictive power is hampered by the lack of information regarding their biological producers. Branched GDGTs have been detected in just a few species of the Acidobacteria but there are strong indications that other bacterial phyla also contribute to the pool of brGDGTs in the environment. Here, we report for the first time the production of potential brGDGT intermediates in Bacillota species. This study demonstrates that brGDGTs likely occur much more widespread in the bacterial domain than previously thought and opens a new chapter both in the understanding of the function of these membrane lipids and their use in paleoclimatology.

## Introduction

Bacterial membrane lipids are generally characterized by fatty acids linked through ester bonds to a glycerol-3-phosphate (G3P) backbone, while archaeal membrane lipids are formed by isoprenoid alkyl moieties linked through ether bonds to glycerol-1-phosphate. In addition, a wide variety of archaea contain predominantly membrane-spanning lipids (MSL), so-called glycerol dialkyl glycerol tetraethers (GDGTs), constituting a monolayer (1). In contrast, bacterial membranes are commonly built up as lipid bilayers. However, there are exceptions to this rule as some bacterial membrane lipids are also formed from alkyl chains spanning across the membrane and linked to the G3P backbone through either ester or ether bonds (2). When these lipids contain four ether bonds and methylations on the alkyl chains (creating branching) they are generally known as branched GDGTs (brGDGTs) (3, 4).

BrGDGTs have been detected in diverse environments including marine sediments (5), soils (6, 7), peats (8), lakes (9, 10), rivers (11, 12), and hydrothermal settings (13, 14). BrGDGTs in ancient marine and lacustrine sediments and loess deposits have been widely used as paleoenvironmental proxies to reconstruct mean annual air temperature, pH, and soil input (15–17). A severe complication in the application of these proxies is that the biological sources of brGDGTs are still poorly understood, hampering proxy validation studies.

In a molecular ecological study of a peat deposit, Acidobacteria were suggested as a potential biological source of environmental brGDGTs (18). Examination of 17 cultures of members of subdivisions (SDs) 1 and 3 of the Acidobacteria showed a high abundance (20-40% of the total fatty acids) of 13,16-dimethyl octacosanedioic acid, hereafter referred to as *iso*-diabolic acid (*iso*-DA; likely resulting from the tail-to-tail coupling of two *iso*-C_15:0_ fatty acids) after acid hydrolysis of the whole cell material (19). *Iso*-DA bears a structural resemblance to the alkyl chains of brGDGTs, and hence its presence suggested the potential of Acidobacteria to synthesize brGDGTs. Indeed, two of the studied Acidobacteria, *Edaphobacter aggregans* Wbg-1 and *Acidobacteriaceae* bacterium A2-4c, contained small amounts of *iso*-DA glycerol ether and a brGDGT comprised of two ether-bound *iso*-DAs. In a subsequent study of *E. aggregans,* it was shown that oxygen limitation can increase the production of brGDGTs (20). Conversely, in most SD4 acidobacterial cultures, *iso*-DA glycerol ether is much more abundant, but brGDGTs were not detected (21, 22). A survey of 46 acidobacterial strains covering 7 different SDs revealed the presence of *iso*-DA in Acidobacteria of the SDs 1, 3, 4 and 6. In some Acidobacteria strains, *iso*-DA containing an additional methyl group at C-5 or C-6 were present (21, 22), lending further support to these groups as biological sources of environmental brGDGTs since these are exactly the positions of additional methylation in environmental brGDGTs (4, 23, 24). More recently, *Ca.* Solibacter usitatus Ellin6076, a member of the Acidobacteria SD3, has been shown to produce a variety of brGDGTs, including C_5_-methylated and cyclopentane-containing brGDGTs in high abundance. A recent study (25) reported the identification of *iso*-DA-based diglycerol tetraesters and their C-6 methyl derivatives in the Acidobacterium *Ca.* Koribacter versatilis Ellin345.

Constraining the effects of environmental and physiological factors on the production of brGDGTs using bacterial cultures is of key relevance to further strengthen the information inferred by using brGDGT-based paleoproxies, which are now only based on empirical calibrations (6, 26–28). Indeed, recent culture work with acidobacterial species has shed further light on the production of brGDGTs. *S. usitatus* Ellin6076 was found to adjust the degree of methylation of brGDGTs according to growth temperature, and the degree of cyclization of brGDGTs is influenced by temperature, pH and oxygen availability (29, 30). In a similar way, the degree of methylation at C-6 of the membrane-spanning *iso*-DA tetraesters decreases with growth temperature in *Ca.* K. versatilis Ellin345 (25), in much the same way as brGDGTs in the environment (6, 26).

A complication in the proxy validation of brGDGTs is that there is circumstantial evidence that they can be produced by bacterial groups other than Acidobacteria, specifically (facultative) anaerobic heterotrophic bacteria. This is based on three different lines of observations. Firstly, culture studies have shown that *iso-*DA is not only biosynthesized by Acidobacteria but also by other bacteria (31–33). Secondly, the brGDGT abundance in the environment is often correlated with the abundance of other bacterial groups (34–37), although this remains an empirical observation. Indeed, previous studies have observed high brGDGT concentrations in the anoxic part of the water column of lakes, where Acidobacteria were almost absent as revealed by the 16S rRNA gene distribution (36, 38), and also by brGDGT production in anoxic marine sediments (39), and in peatland soils (40). Thirdly, the genes coding for the enzymes involved in the coupling of the alkyl chains, i.e., Mss, membrane spanning lipid synthase, and of the ether bond formation, i.e., Ger, glycerol ester reductase (Fig. 1), likely involved in the bacterial biosynthesis of membrane-spanning ether lipids, are also found in the genomes of bacterial groups other than Acidobacteria (33). In this regard, a screening of the presence of these biosynthetic genes in the euxinic waters of the Black Sea also pointed to non-Acidobacterial, anaerobic bacterial taxa as potential biological producers of brGDGTs (41).

**Figure 1.**
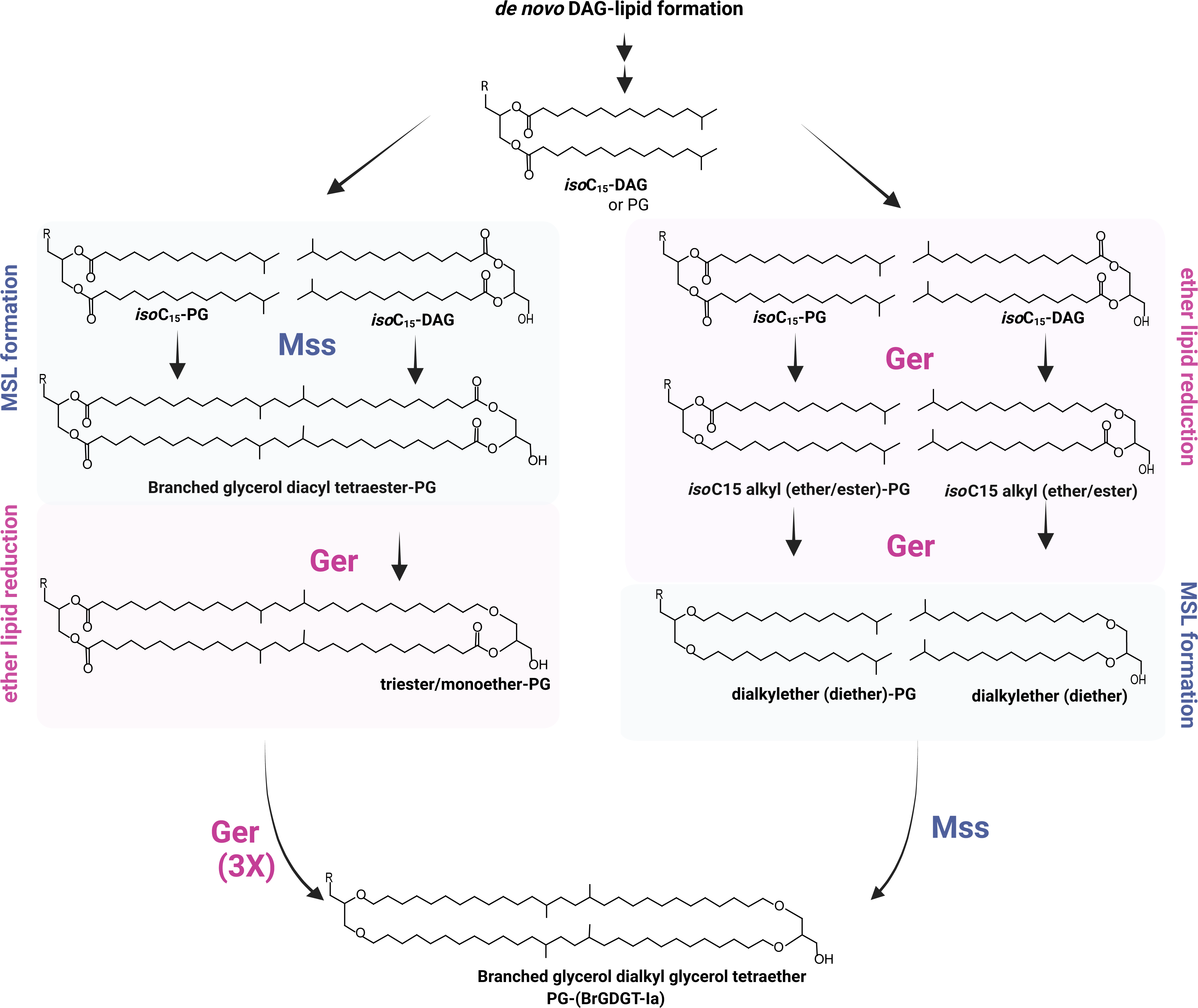
Hypothetical biosynthetic pathway for brGDGTs via Mss and Ger enzymes. This proposed pathway suggests that diacylglycerol (DAG), specifically i*so-*C_15_-DAG, is a biosynthetic precursor by either the membrane spanning lipid synthase (Mss; Left) or glycerol ester reductase (Ger; Right), the order of which (if there is any) remains unknown. The Mss condenses *iso-*C_15:0_-DAG monomers into branched glycerol diacyl tetraesters compounds which subsequently can be reduced by Ger at the *sn*-1 and *sn*-2 position converting the tetraester (intermediate) into a fully reduced brGDGT-Ia (bottom). Ger can utilize *iso*-C_15:0-_DAG to produce monoethers followed by diethers intermediates, which are then further processed by Mss to produce brGDGTs. R=H

We previously reported the presence of the *mss* and *ger* biosynthetic genes and the presence of *iso*-DA and glycerol ethers in cultures of the bacterial phylum Bacillota (formerly known as Firmicutes), of the Class Clostridia (33). *Bacillota* species are important inhabitants in soils and other environmental systems (42). However, the production of brGDGTs by this phylum has never been demonstrated. Here, we report for the first time the identification of putative biosynthetic brGDGT intermediates in the Bacillota phylum, which provides further support for the hypothesis that brGDGTs are likely also biosynthesized outside the phylum Acidobacteria. We also present a biosynthetic pathway leading to the formation of these membrane lipids.

## 2. Results and Discussion

### 2.1 Genomic potential for brGDGT production within the Bacillota

We first searched for the co-occurrence of the two essential genes for brGDGT biosynthesis: the membrane spanning synthase (*mss*), and the glycerol ester reductase (*ger*) (Table 1, Supplementary Table 1) in genomes of the phylum Bacillota. We observed the genomic co-occurrence of these genes in many members, distributed across 6 classes and 54 families (Table 2; Supplementary Tables 2-4). From these, we selected 7 strains to study potential biosynthesis of brGDGTs. Firstly, five members of Class Clostridia which had previously been found to produce *iso*-DA (33, 43). Two additional strains from the Class Tissierellia were chosen based on their environmental significance (Supplementary Table 2), and on the availability of the strains in culture collections: *Keratinibaculum paraultenense* (family Tepidimicrobiaceae), a thermophilic, spore-forming, anaerobic bacterium isolated from grassy marshlands (44), and *Sporoanaerobacter acetigenes* LUP 33T (family Sporanaerobacteraceae), a strictly anaerobic, moderately thermophilic, originally isolated from anaerobic sludge (45). Both strains are phylogenetically closely related to the Class Clostridia (Table 1). All strains were grown to stationary phase at optimal growth conditions.

**Table 1.** Overview of lipids detected in Bacillota strains which harbor both the membrane spanning lipid synthase (*mss*) and glycerol ester reductase (*ger*) coding genes. Bligh and Dyer extracts (BDE) of these strains were analyzed by UHPLC-HRMS^n^. Acid-hydrolyzed biomas**s** was analyzed both by UHPLC-HRMS^n^ and GC-MS (see text for details). See Supplementary Table 1 for further details on the taxonomy, metabolism, and core lipid analysis.

|  |  |  | UHPLC-HRMS<sup>n</sup> detection |  |  |  | GC-MS detection |
| --- | --- | --- | --- | --- | --- | --- | --- |
| Strain | Genomic search<sup>a</sup> |  | BDE |  | Acid-hydrolyzed biomass |  | Acid-hydrolyzed biomass |
|  | Mss | Ger | brGDGTs | brGDGT ester analogues | brGDGTs | brGDGT ester analogues | *Iso*-diabolic acids & related compounds<sup>d</sup> |
| *F. ferrireducens*<br>DSM 21121 | ✓ | ✓ | -<sup>b</sup> | - | - | - | *iso*-DA (13,16-diMe C28:0 diFA), 13-Me C28:0 diFA<sup>c</sup> |
| *T. wiegelii*<br>DSM 10319 | ✓ | ✓ | - | - | - | - | *iso*-DA<sup>c</sup> |
| *C. subterraneus*<br>DSM 15729 | ✓ | ✓ | - | - | - | - | *iso*-DA; 30-OH-13,16-diMe C30:0 FA<sup>c</sup> |
| *M. thermoacetica*<br>DSM 12797 | ✓ | ✓ | - | - | - | - | *iso*-DA; N-glyceroyl alkylamines with C30–32 *iso*-DA like structures<sup>c</sup> |
| *C. owensensis*<br>DSM 13100 | ✓ | ✓ | - | - | - | - | Extended *iso*-DA (15,18-diMe C32:0 diFA), 15-Me C32:0 diFA<sup>c</sup> |
| *K. paraultunense*<br>DSM 26752 | ✓ | ✓ | - | ✓ | - | - | *iso*-DA<br>*iso*-DA with 1 vinyl ether (13,16-diMe-28-hydroxyoctacos-27-enoic acid)<br>*iso*-DA with 2 vinyl ether (13,16-diMe-1,28-dihydroxyoctacos-1,27-diene) |
| *S. acetigenes*<br>DSM 13106 | ✓ | ✓ | - | ✓ | - | - | *iso*-DA<br>*iso*-DA with 1 vinyl ether (13,16-diMe-28-hydroxyoctacos-27-enoic acid)<br>*iso*-DA with 2 vinyl ether (13,16-diMe-1,28-dihydroxyoctacos-1,27-diene) |

<sup>a</sup> protein homology search as described in (33) and this study.
<sup>b</sup> refers to lack of detection.
<sup>c</sup> previously reported by (33) and (43).
<sup>d</sup> lipids are listed in decreasing order of abundance (%). diMe= dimethyl, diFA = two carboxylic acid moieties on chain.

**Table 2.**
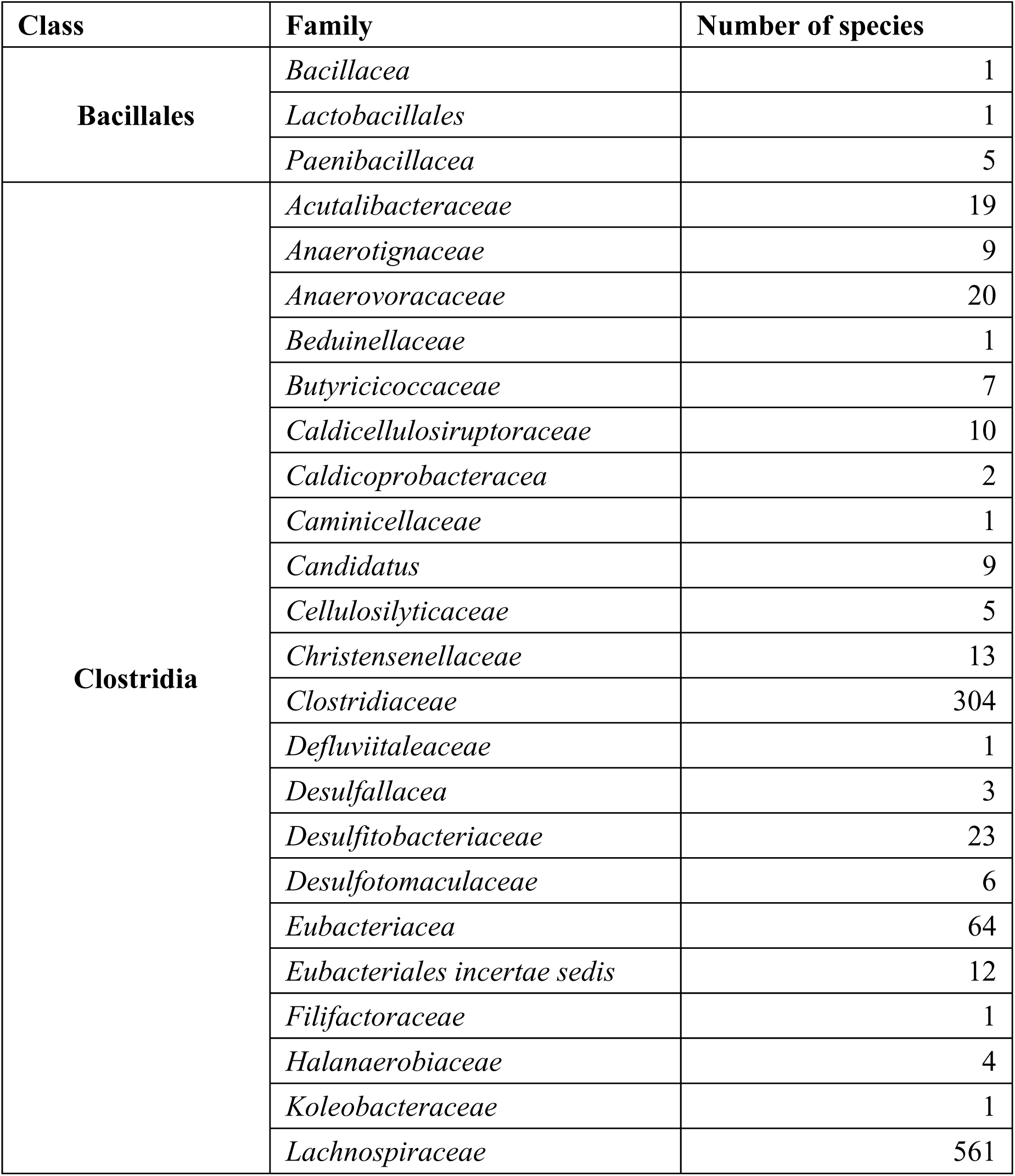
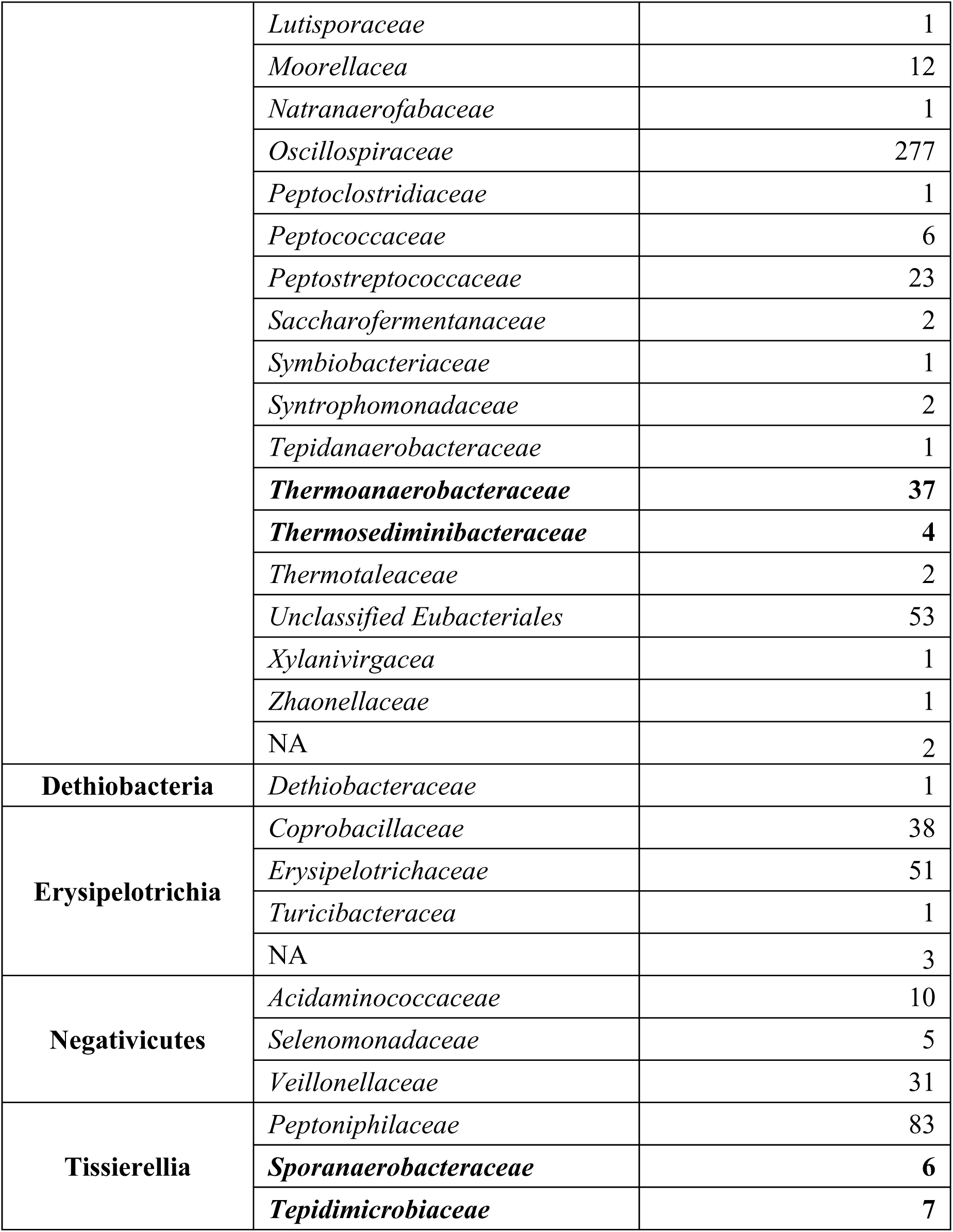
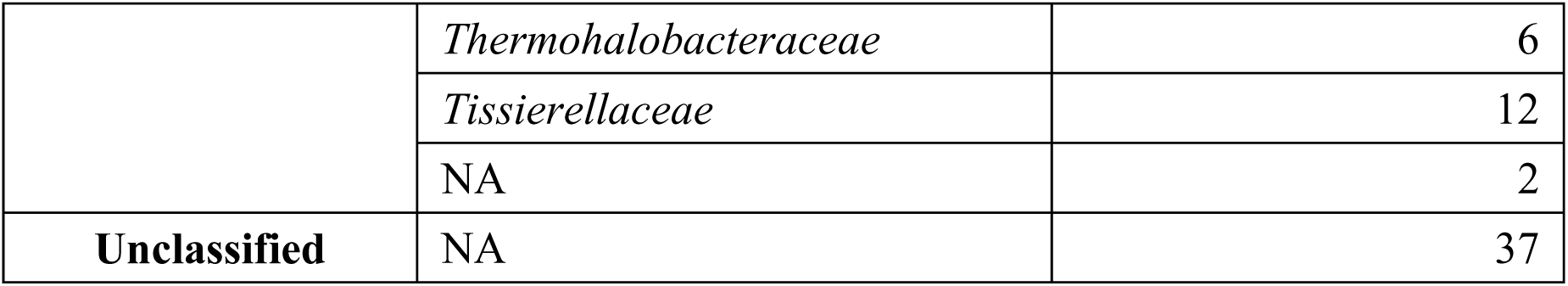
Summary of the number of species distributed across six bacterial class and fifty-six families with co-occurrence of the membrane spanning lipid synthase (Mss) and glycerol ester reductase (Ger) coding gene homologs in their genomes (see material and methods for search criteria). The four families of strains examined in this study are indicated in bold.

### 2.2 Analysis of brGDGTs and related lipids

Several analytical approaches have been reported to screen for brGDGTs or their building blocks. *Iso*-DA has been reported to only be released from acidobacterial biomass after base or acid hydrolysis of cell material, or hydrolysis of the residue of the cell material after Bligh and Dyer lipid extraction (19). On the other hand, both brGDGTs and *iso*-DA-based diglycerol tetraesters have been detected in non-hydrolyzed lipid extracts of environmental samples (46), and in isolated strains as indicated above. Here, we analyzed the 7 selected strains by gas chromatography-mass spectrometry (GC-MS) for hydrolysis-derived *iso*-DA and related compounds and by ultra-high pressure liquid chromatography high resolution multi-stage mass spectrometry (UHPLC-HRMS^n^) to screen for both non-hydrolyzed (intact) brGDGTs and hydrolysis-derived brGDGTs.

#### 2.2.1 Acid hydrolysis-derived lipids

The acid hydrolysis-derived lipids for the 4 strains of the Class Clostridia and one of the Class Bacillota incertae sedis (*Caldicellulosiruporales*) have previously been reported (33, 43), and they all contain *iso*-DA or closely related >C_30_ diacids. For *K. paraultunense* the main acid hydrolysis-derived lipids were *iso*-C_15_ fatty acid (FA) and *iso*-C_15_-dimethylacetal (DMA; formed when alkenyl or also called vinyl ethers are acid-hydrolyzed (47), representing 54% and a 16% of the total lipids detected, respectively, and the membrane-spanning *iso*-DA (Fig. 2A), which comprised 18% of the total (Supplementary Table 5). In the case of *S. acetigenes,* the acid hydrolysis-derived lipids consisted predominantly of *iso*-C_15_ FA and *iso*-C_15_-DMA (representing 37% and 16% of the total, respectively) and *iso*-DA (25%) (Supplementary Table 6).

**Figure 2.**
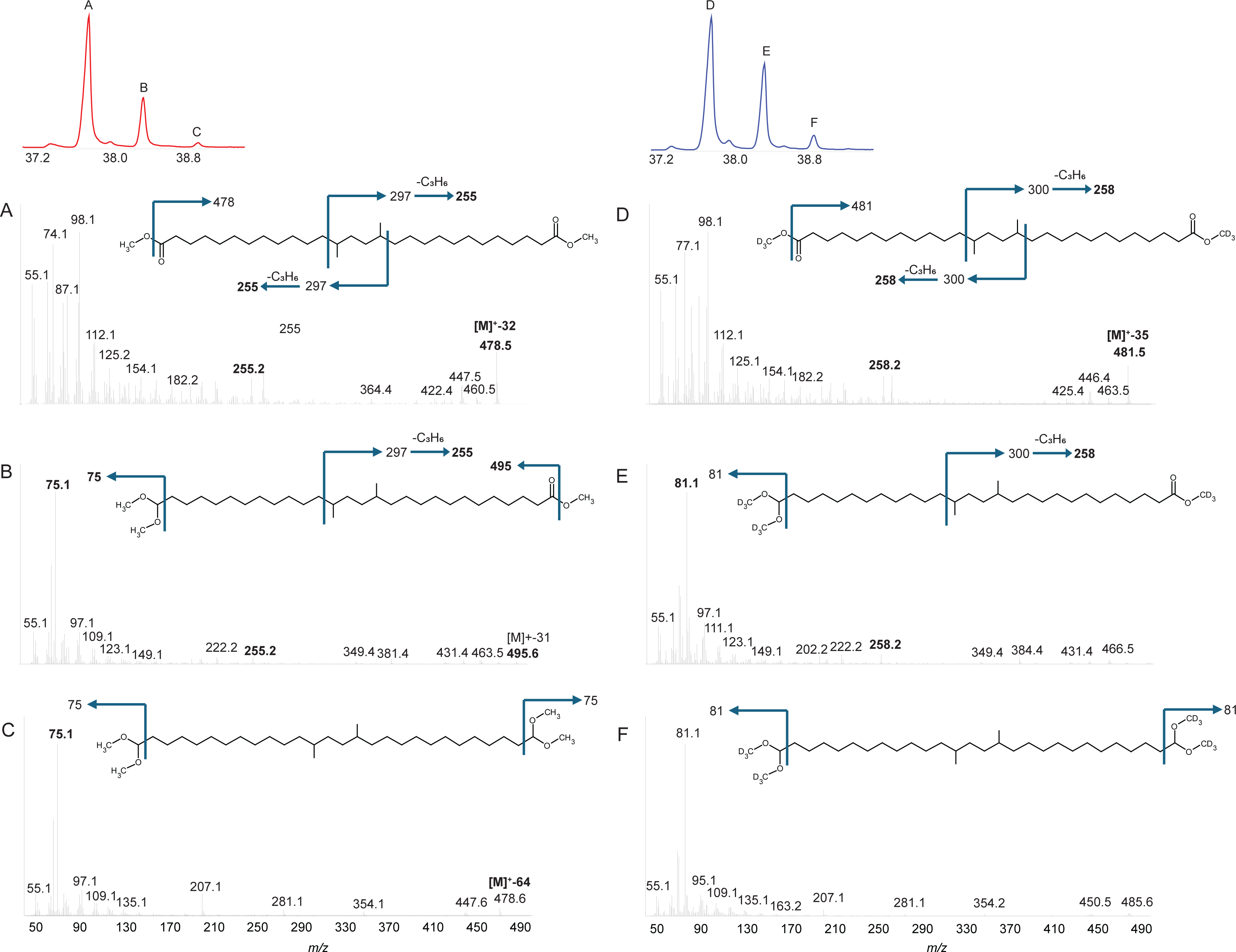
EI (70 eV) mass spectra obtained by analysis with gas chromatography-mass spectrometry (GC-MS) of high-molecular weight lipids detected in acid-hydrolyzed biomass of *K. paraultunense*. A) *iso*-diabolic acid (13,16-dimethyl octacosanedioic acid), with both acids in the form of methyl esters; B) 13,16-dimethyl-1-methyl ester,28-dimethylacetal C) 13,16-dimethyl-1,28-bis(dimethylacetal). GC-MS mass spectra of the same three components after deuterated methylation: D) *iso*-diabolic acid methyl ester; E) 13,16-dimethyl-1-methylester,28-dimethylacetal; and F) 13,16-dimethyl-1,28-bis(dimethylacetal). Insert top left: partial TIC with peaks A, B and C indicated. Insert top right: partial TIC with peaks D, E and F indicated. Note that retention times and relative peak intensity are not affected by deuterated methylation.

#### 2.2.2 BrGDGT analogues

BrGDGTs themselves were not detected either in the Bligh and Dyer extracts or in the acid hydrolysis extracts for any of the 7 Bacillota strains, despite the presence of the key biosynthetic genes and the detection of *iso*-DA (or related lipids) in the lipid analysis of acid-hydrolyzed biomass (Table 1; Supplementary Table 1). However, using UHPLC-HRMS^n^ we detected a series of MSLs in the Bligh and Dyer extracts of two of the seven strains: *K. paraultunense* and *S. acetigenes* (Table 1; Supplementary Table 1, Table 3). These compounds eluted earlier than the known brGDGTs during reverse phase separation (41), indicating a more polar character. A series of 3 isomers, present in both species, with an ammoniated molecular ion at *m/z* 1094.952 had the identical elemental composition (C_66_H_128_O_10_N ([M+NH_4_]^+^)) and MS^2^ fragmentation as the *iso*-DA-based diglycerol tetraester (ester analogue of brGDGTs-Ia), reported previously in *T. ethanolicus* (31) and recently reported to occur in highly elevated soils (48), and in the acidobacterium *Ca.* Solibacter usitatus (25). We will subsequently refer to this *iso*-DA-based diglycerol tetraester as brGDGT analogue **1** (Table 3). In addition, we detected a single peak arising from an ammoniated molecular ion at *m/z* 1080.973 (elemental composition C_66_H_130_O_9_N; Table 3), and a single peak arising from ammoniated molecular ion at *m/z* 1066.994 (elemental composition C_66_H_132_O_8_N; Table 3). Based on comparison of their elemental composition and the spectral similarities to brGDGT analogue **1**, we identified these compounds as brGDGT analogues with three ester bonds and one ether bond (brGDGT analogue **2**, Table 3), and with two ester bonds and two ether bonds (brGDGT analogue **3**, Table 3) respectively. A brGDGT analogue with one ester bond and three ether bonds was not detected. Further evidence for these lipids containing ester bonds was their disappearance after acid hydrolysis (Table 1). brGDGT analogues with one and two ester bonds were previously also detected by (48) in soils. A similar series of membrane spanning lipids with increasing number of ether bonds, but with diabolic acid (15,16-dimethyltriacontanedioic acid) instead of *iso*-DA as building blocks were previously identified in various species of the genus *Thermotogota* (49).

**Table 3.**
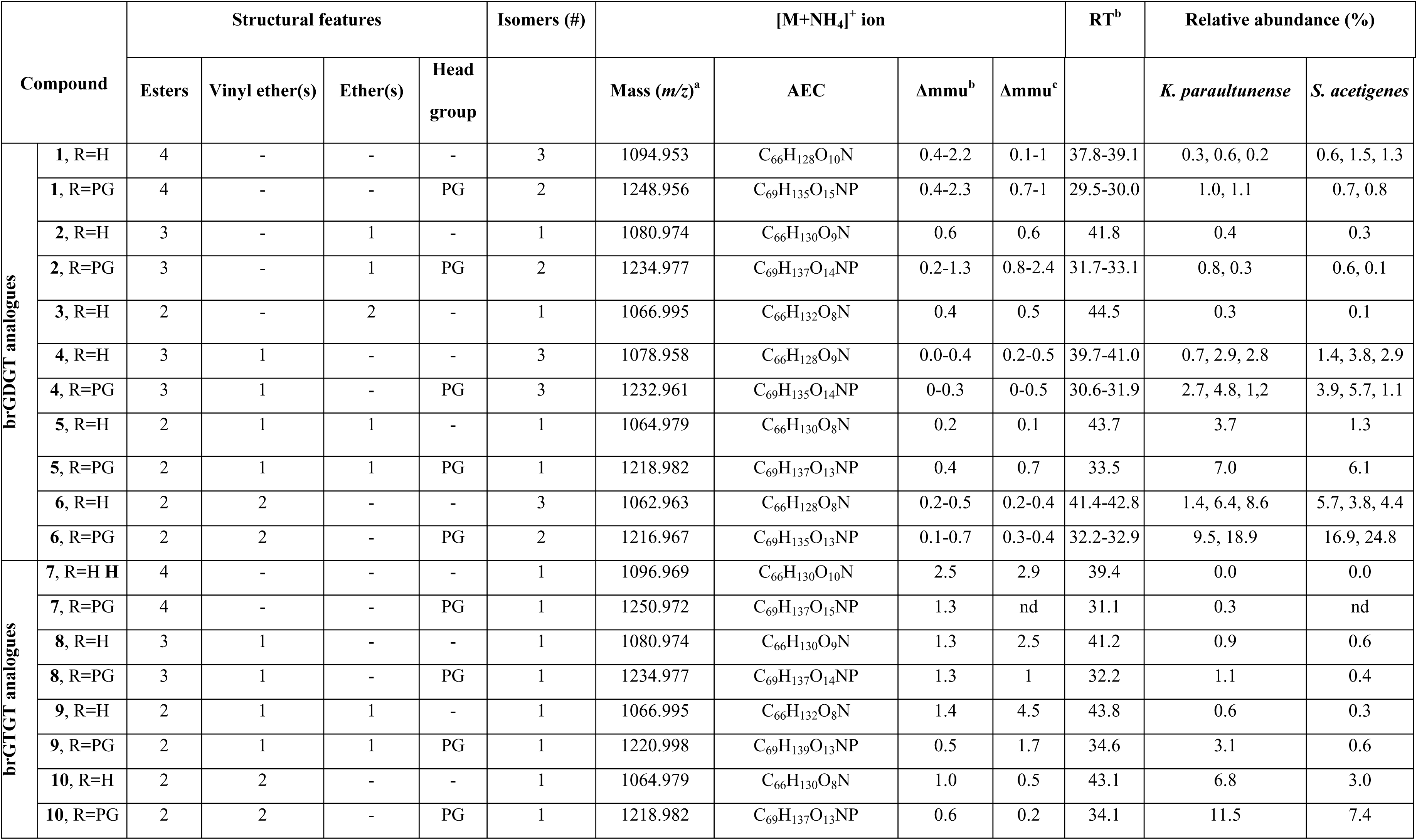
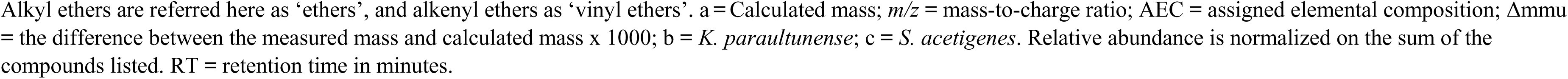
Analogues of brGDGTs and brGTGT detected by UHPLC-HRMS^n^ in *K. paraultunense* and *S. acetigenes* biomass grown at optimal conditions.

| Compound |  | Structural features |  |  |  | Isomers (#) | [M+NH<sub>4</sub>]<sup>+</sup> ion |  |  |  | RT<sup>b</sup> | Relative abundance (%) |  |
| --- | --- | --- | --- | --- | --- | --- | --- | --- | --- | --- | --- | --- | --- |
|  |  | Esters | Vinyl ether(s) | Ether(s) | Head group |  | Mass (m/z)<sup>a</sup> | AEC | Δmmu<sup>b</sup> | Δmmu<sup>c</sup> |  | *K. parautunense* | *S. acetigenes* |
| brGDGT analogues | 1, R=H | 4 | - | - | - | 3 | 1094.953 | C<sub>66</sub>H<sub>128</sub>O<sub>10</sub>N | 0.4-2.2 | 0.1-1 | 37.8-39.1 | 0.3, 0.6, 0.2 | 0.6, 1.5, 1.3 |
|  | 1, R=PG | 4 | - | - | PG | 2 | 1248.956 | C<sub>69</sub>H<sub>135</sub>O<sub>15</sub>NP | 0.4-2.3 | 0.7-1 | 29.5-30.0 | 1.0, 1.1 | 0.7, 0.8 |
|  | 2, R=H | 3 | - | 1 | - | 1 | 1080.974 | C<sub>66</sub>H<sub>130</sub>O<sub>9</sub>N | 0.6 | 0.6 | 41.8 | 0.4 | 0.3 |
|  | 2, R=PG | 3 | - | 1 | PG | 2 | 1234.977 | C<sub>69</sub>H<sub>137</sub>O<sub>14</sub>NP | 0.2-1.3 | 0.8-2.4 | 31.7-33.1 | 0.8, 0.3 | 0.6, 0.1 |
|  | 3, R=H | 2 | - | 2 | - | 1 | 1066.995 | C<sub>66</sub>H<sub>132</sub>O<sub>8</sub>N | 0.4 | 0.5 | 44.5 | 0.3 | 0.1 |
|  | 4, R=H | 3 | 1 | - | - | 3 | 1078.958 | C<sub>66</sub>H<sub>128</sub>O<sub>9</sub>N | 0.0-0.4 | 0.2-0.5 | 39.7-41.0 | 0.7, 2.9, 2.8 | 1.4, 3.8, 2.9 |
|  | 4, R=PG | 3 | 1 | - | PG | 3 | 1232.961 | C<sub>69</sub>H<sub>135</sub>O<sub>14</sub>NP | 0-0.3 | 0-0.5 | 30.6-31.9 | 2.7, 4.8, 1.2 | 3.9, 5.7, 1.1 |
|  | 5, R=H | 2 | 1 | 1 | - | 1 | 1064.979 | C<sub>66</sub>H<sub>130</sub>O<sub>8</sub>N | 0.2 | 0.1 | 43.7 | 3.7 | 1.3 |
|  | 5, R=PG | 2 | 1 | 1 | PG | 1 | 1218.982 | C<sub>69</sub>H<sub>137</sub>O<sub>13</sub>NP | 0.4 | 0.7 | 33.5 | 7.0 | 6.1 |
|  | 6, R=H | 2 | 2 | - | - | 3 | 1062.963 | C<sub>66</sub>H<sub>128</sub>O<sub>8</sub>N | 0.2-0.5 | 0.2-0.4 | 41.4-42.8 | 1.4, 6.4, 8.6 | 5.7, 3.8, 4.4 |
|  | 6, R=PG | 2 | 2 | - | PG | 2 | 1216.967 | C<sub>69</sub>H<sub>135</sub>O<sub>13</sub>NP | 0.1-0.7 | 0.3-0.4 | 32.2-32.9 | 9.5, 18.9 | 16.9, 24.8 |
| brGTGT analogues | 7, R=H H | 4 | - | - | - | 1 | 1096.969 | C<sub>66</sub>H<sub>130</sub>O<sub>10</sub>N | 2.5 | 2.9 | 39.4 | 0.0 | 0.0 |
|  | 7, R=PG | 4 | - | - | PG | 1 | 1250.972 | C<sub>69</sub>H<sub>137</sub>O<sub>15</sub>NP | 1.3 | nd | 31.1 | 0.3 | nd |
|  | 8, R=H | 3 | 1 | - | - | 1 | 1080.974 | C<sub>66</sub>H<sub>130</sub>O<sub>9</sub>N | 1.3 | 2.5 | 41.2 | 0.9 | 0.6 |
|  | 8, R=PG | 3 | 1 | - | PG | 1 | 1234.977 | C<sub>69</sub>H<sub>137</sub>O<sub>14</sub>NP | 1.3 | 1 | 32.2 | 1.1 | 0.4 |
|  | 9, R=H | 2 | 1 | 1 | - | 1 | 1066.995 | C<sub>66</sub>H<sub>132</sub>O<sub>8</sub>N | 1.4 | 4.5 | 43.8 | 0.6 | 0.3 |
|  | 9, R=PG | 2 | 1 | 1 | PG | 1 | 1220.998 | C<sub>69</sub>H<sub>139</sub>O<sub>13</sub>NP | 0.5 | 1.7 | 34.6 | 3.1 | 0.6 |
|  | 10, R=H | 2 | 2 | - | - | 1 | 1064.979 | C<sub>66</sub>H<sub>130</sub>O<sub>8</sub>N | 1.0 | 0.5 | 43.1 | 6.8 | 3.0 |
|  | 10, R=PG | 2 | 2 | - | PG | 1 | 1218.982 | C<sub>69</sub>H<sub>137</sub>O<sub>13</sub>NP | 0.6 | 0.2 | 34.1 | 11.5 | 7.4 |

Alkyl ethers are referred here as ‘ethers’, and alkenyl ethers as ‘vinyl ethers’. a = Calculated mass; $m/z$ = mass-to-charge ratio; AEC = assigned elemental composition; $\Delta m$ = the difference between the measured mass and calculated mass x 1000; b = *K. paraultunense*; c = *S. acetigenes*. Relative abundance is normalized on the sum of the compounds listed. RT = retention time in minutes.

Several other components exhibited similar masses to the brGDGT analogues **1-3,** but had accurate masses indicative of the presence of double bond equivalents (DBE). During GC-MS analysis of the acid-hydrolyzed biomass, several DMAs were detected, which are formed upon acid hydrolysis of vinyl ether lipids (as found in plasmalogen lipids) as determined by the presence of a dominant ion at *m/z* 75. These included *iso*-C_15_ DMA (Table 1) as well as high-molecular weight lipids. These likely represent the dimethylacetalized form of compounds with a 13,16-dimethyloctacosanyl chain, structurally analogous to *iso*-DA (Fig. 2A), but containing one vinyl ether bond (13,16-diMe-28-hydroxyoctacos-27-enoic acid; Fig. 2B) or two vinyl ether bonds (13,16-diMe-1,28-dihydroxyoctacos-1,27-diene; Fig 3C). To confirm whether the *m/z* 75 fragment was indeed derived from DMA moieties, an aliquot of the biomass was hydrolyzed with deuterated methanol. The fragment at *m/z* 75 was now detected at *m/z* 81 (a shift of 6 Da), confirming that the fragment contained two methoxy groups.

**Figure 3.**
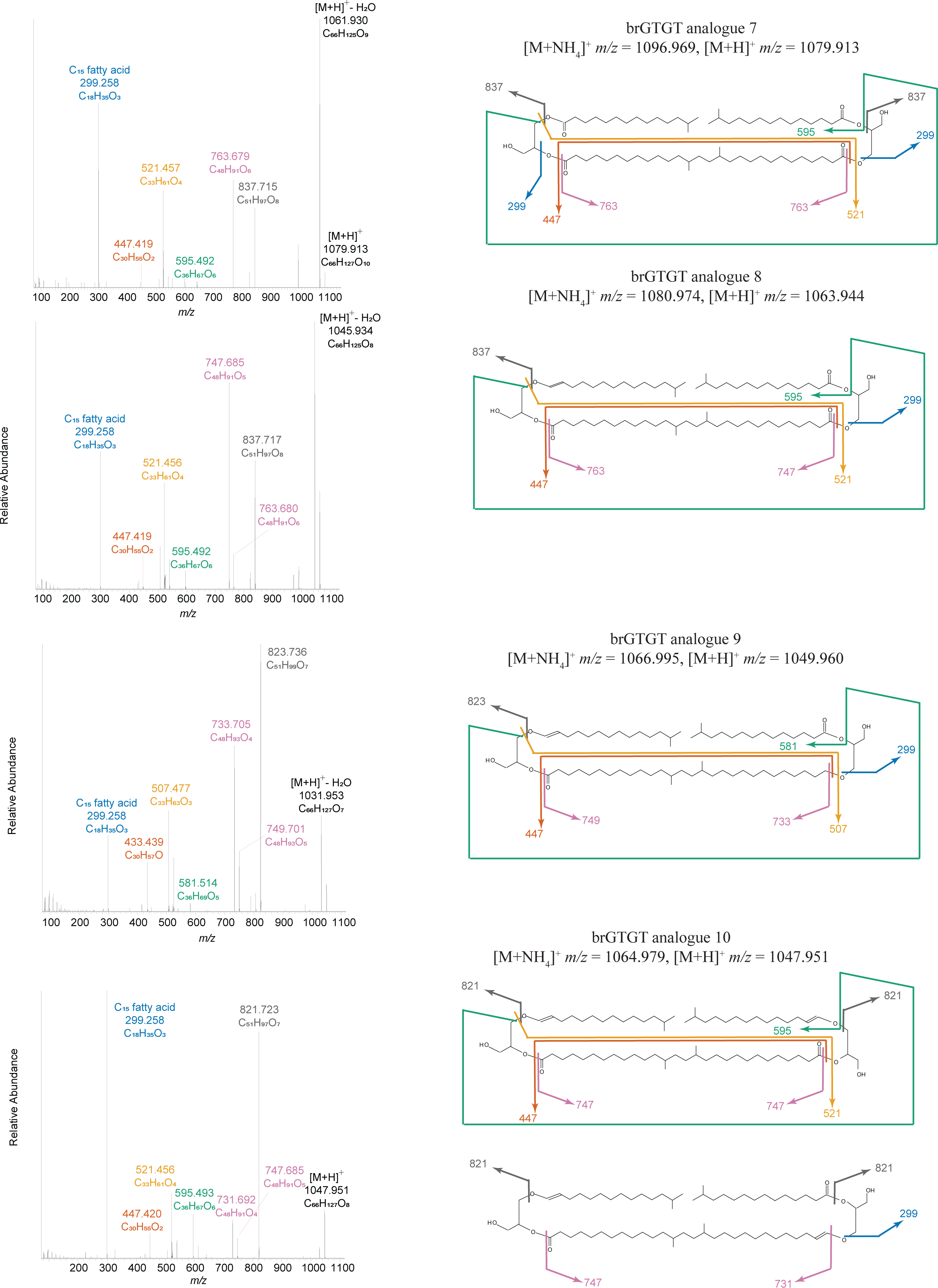
MS^2^ spectra of brGTGT analogues **7-10** obtained by UHPLC-HRMS^n^ analysis of lipid extract from *K. paraultunense*. brGTGT analogue **10** exhibited a complex spectrum, suspected to represent a mixture of two structural isomers.

Based on this GC-MS analysis, the brGDGT analogues with DBEs, observed during UHPLC-MS^n^ analysis, were assumed to contain vinyl ether bonds. Three isomers with an [M+NH_4_]^+^ at *m/z* 1078.958 (C_66_H_128_O_9_N) were assigned as brGDGT analogues with three ester bonds and one vinyl ether bond (brGDGT analogue **4,** Table 3). A single component with [M+NH_4_]^+^ at *m/z* 1064.979 (C_66_H_130_O_8_N) was assigned as a brGDGT analogue with two ester bonds, one ether bond and one vinyl ether bond (brGDGT analogue **5**, Table 3). While three isomers with *m/z* values of 1062.963 (C_66_H_128_O_8_N) were assigned as brGDGT analogues with two ester bonds and two vinyl ether bonds (brGDGT analogue **6**, Table 3).

#### 2.2.3. brGTGT analogues

In both *K. paraultunense* and *S. acetigenes* four additional components with [M+NH_4_]^+^ ions with *m/z* 1096.969 (C_66_H_130_O_10_N), *m/z* 1080.973 (C_66_H_130_O_9_N), 1066.994 (C_66_H_132_O_8_N) and 1064.979 (C_66_H_130_O_8_N), were detected (Table 3).

While these were identical in mass to several of the brGDGT analogues described above (Table 3), they exhibited different retention times and differences in their MS^2^ spectra. These MS^2^ spectra are highly comparable with the mass spectrum of a branched glycerol trialkyl glycerol tetraether (brGTGT; one side of lipid is ‘open’) described by (20) as well as the mass spectrum of isoprenoid GTGTs described by (50). Their MS^2^ spectra (Fig. 3) all exhibited a major fragment at *m/z* 299.258, indicative of a glycerol- C_15_ fatty acid moiety, as well as a number of dominant ions in the *m/z* 700-830 range, which are generally minimal in the MS^2^ of brGDGTs (51) and in the brGDGT analogues described above. The elemental compositions of these fragments indicate they arise from loss of a C_15_ moiety (in case of the C_51_ fragments) or a C_15_-glycerol moiety (in case of the C_48_ fragments). In addition, three of these components contained 1 or 2 DBE, which together with the above-described detection of both iso-C_15_ DMA and long-chain DMA products after GC analysis, indicate the presence of vinyl ethers, similar to the vinyl ether-containing brGDGT analogues described above. Hence, these three components were assigned as brGTGT analogues: *m/z* 1096.969 containing four ester bonds (brGTGT analogue **7**, Table 3), *m/z* 1080.973 with three ester bonds and one vinyl ether bond (brGTGT analogue **8**, Table 3), *m/z* 1066.994 with two ester bonds, one ether bond and one vinyl ether bond (brGTGT analogue **9**, Table 3) and *m/z* 1064.979 with two ester bonds and two vinyl ether bonds (brGTGT analogue **10**, Table 3).

Further inspection of the brGTGT analogue fragmentation spectra allowed for assignment of the distribution of ether, vinyl-ether or ester bonds within the molecule. The proposed fragmentation pathways are shown in Fig. 3. BrGTGT analogue **7 (***m/z* 1096.969, four ester bonds, Table 3), exhibited an MS^2^ spectrum indicating that **7** comprised two C_15_ fatty acids and a membrane spanning *iso*-DA (Fig. 3). The brGTGT **8** analogue (*m/z* 1080.974, three ester bonds, Table 3), exhibited an MS^2^ spectrum indicating that **8** comprised a membrane spanning *iso*-DA, a C_15_ fatty acid and a C_15_ vinyl ether moiety (Fig. 3). This composition suggests that one part of this lipid is in the form of a plasmalogen, in which the *sn*-1 position of the glycerol is bound to a 1-*O*-alk-1′-enyl moiety (i.e. a vinyl ether), whereas the *sn*-2 position is an acyl ester. BrGTGT analogue **9** (*m/z* 1066.995, two ester bonds, Table 3) exhibited an MS^2^ spectrum with several differences to those of brGTGT analogues **7** and **8**. These differences indicated that this does not comprise an esterified membrane spanning *iso*-DA, but instead a membrane spanning 13,16-dimethyl octacosanyl chain with an acid at one end and an ether bond at the other, with the ‘open’ side comprising a C_15_ fatty acid and a C_15_ vinyl ether moiety (Fig. 3). Finally, brGTGT analogue **10** (*m/z* 1064.979, two ester bonds, Table 3) exhibited a more complex spectrum, with variability in the relative proportion of the fragment ions across the peak, which was broader than that of the other three brGTGT analogues. Indeed, we have tentatively assigned it as being mix of two structural isomers (Fig. 3). The first contains an esterified membrane spanning *iso*-DA and two C_15_ vinyl ether moieties, while the second contains a 13,16-dimethyl octacosanyl chain with both an acid and a vinyl ether bond, as well as a C_15_ fatty acid and a C_15_ vinyl ether moiety. We assume that in both isomers the vinyl ether bonds would be at the *sn*-1 position of the glycerols and that the ester bonds would be at the *sn*-2 position, as is the case for plasmalogen lipids (47). Similar lipids have been described in the Bacillota genus *Butyrivibrio* (52) but then with DA as the membrane spanning moiety and with two C_16_ vinyl ether moieties.

While we present both of the proposed structural isomers of brGTGT analogue **10** in Figure 3, we have only presented one of them in Figure 4. That is the isomer which contains a 13,16-dimethyl octacosanyl chain with an acid at the *sn*-2 position of one glycerol and a vinyl ether bond at the *sn*-1 position of the other glycerol. This conformation is in line with how we have drawn brGDGT analogues **1**-**6** (Fig. 4), *i.e*., with an anti-parallel configuration (53). It should be noted, however, that we have presented them in this form as it is most routinely described in the literature and that it is actually impossible to assign either an anti-parallel or parallel configuration from their UHPLC-HRMS^n^ spectra to any of the brGDGT analogues described here.

**Figure 4.**
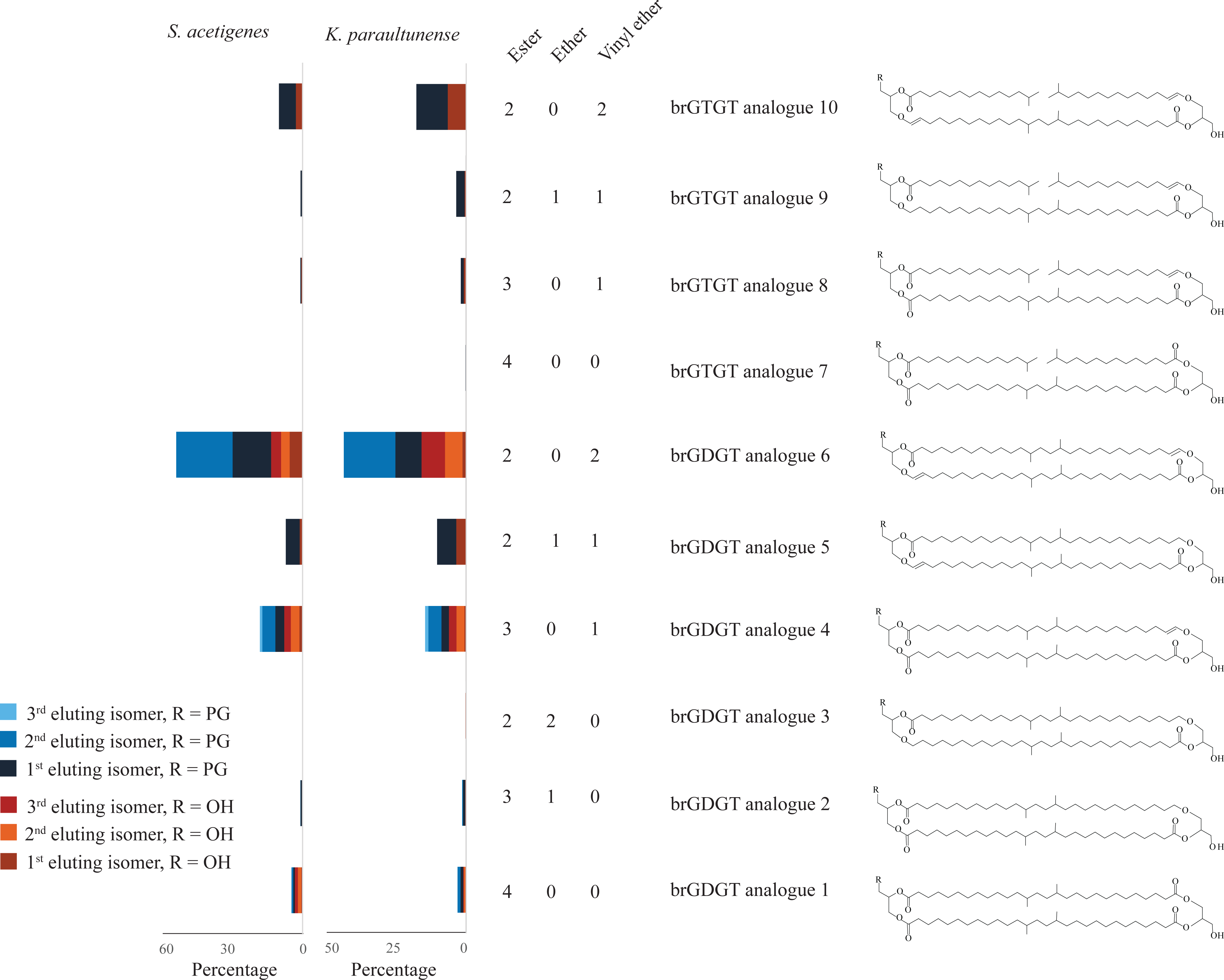
The relative proportions of the various brGDGT analogues (**1–6**) and brGTGT analogues (**7–10**) and isomers thereof and tentative examples of structures. It should be noted that it would be expected that the lipids with and without polar head groups would have considerable differences in their degree of ionization efficiency during LC/MS analysis. Hence, the relative abundances shown here, based on peak areas (in response units), do not necessarily reflect their actual abundance. However, this quasi-quantification allows for comparison between the two samples. As explained in the text, we have not determined whether the components have an anti-parallel or a parallel configuration, for simplicity have presented them all with an anti-parallel configuration.

#### 2.2.5 Intact polar brGDGT and brGTGT analogues

For each of the brGDGT and brGTGT analogues, a corresponding intact polar lipid with one phosphoglycerol (PG) head group was identified in the Bligh and Dyer extracts of *K. paraultunense* and *S. acetigenes* (Table 3). The intact polar brGDGT and brGTGT analogues were identified based on their accurate masses (Table 3) and the characteristic neutral loss in MS^2^ of 189.04 Da (C_3_H_12_O_6_NP), associated with loss from an ammoniated ion of a phosphoglycerol moiety and NH_3_. No other head groups were detected as part of the brGDGT or brGTGT analogues. An exception to this was the brGDGT analogue **3** (*m/z* 1066.994; Table 3), which was not detected in intact polar form, possibly due to low abundance.

#### 2.2.6 Distribution of brGDGT and brGTGT analogues

For both species grown under optimum growth conditions, the most abundant brGDGT analogue, both with and without a PG head group, was brGDGT analogue **6**, containing two ester bonds and two vinyl ether bonds (Fig. 4 and Table 3).Similarly, the most abundant brGTGT analogue, both with and without a PG head group was brGTGT analogue **9**, again with two ester bonds and two vinyl ether bonds (Fig. 4 and Table 3).

It is notable that the brGDGT analogues detected in both *K. paraultunense* and *S. acetigenes* exhibited variable numbers of isomers as detected by UHPLC-HRMS^n^ (Fig. 4 and Table 3). For example, three isomers of brGDGT analogue **4** were detected with a PG head group (*m/z* 1232.962) and a further three without (*m/z* 1078.958). Conversely, only one isomer of brGDGT analogue **5** was detected with a PG head group (*m/z* 1218.982) and one without (*m/z* 1064.979). There seems to be a correlation between the number of isomers and the presence of certain bonds. Those brGDGT analogues with one or more ether bonds appear to have fewer isomers than those with none. This was not the case for the brGTGT analogues, as for each one, only one isomer was detected, both with and without a PG head group (Fig. 4 and Table 3). It is interesting to note that other studies which have reported membrane spanning lipids with ester bonds have also reported more isomers for components with no ether bonds, than for those with (25, 48, 49).

### 2.3 Effect of growth conditions on the lipid composition

After identification of brGDGT and brGTGT analogues in *K. paraultunense* and *S. acetigenes*, we examined the effect of temperature and oxygen concentration on the production of these lipids. In the environment, temperature exerts a major role in brGDGT distribution which has led to the development of brGDGT proxies for the reconstruction of past temperatures (6, 7). Oxygen has been shown to affect the production of brGDGTs in aerobic members of the Acidobacteria (19, 20, 29). Considering that the two Bacillota strains *K. paraultunense* and *S. acetigenes* are strictly anaerobic, oxygen exposure is a particularly pertinent variable to examine in relation to the production of brGDGT analogues.

To quantify changes in the brGDGT analogue lipids, we carried out acid-hydrolysis and quantified the resulting products using GC-MS. It is to be expected that the brGDGT analogues with and without polar head groups have differences in their degree of ionization efficiency during LC-MS analysis. Therefore, examination of the acid hydrolysis-derived products of the lipids by GC-MS allows for more accurate quantification. To this end, we determined changes in the percent of MSLs (i.e., sum of all 13,16-dimethyloctacosanyl-based compounds, both ester and ether bound; Table 4). We also examined the percent of hydrolysis-derived lipids with an ether bond (Table 4).

**Table 4.** Percentage of hydrolysis-derived lipids that are membrane spanning (MSLs) and containing an ether bond.

| Species |  | Temperature (°C) |  |  |  |  |  | Oxygen level (%)** |  |  |
| --- | --- | --- | --- | --- | --- | --- | --- | --- | --- | --- |
|  |  | 27 | 37* | 47 | 45 | 55* | 60 | 1 | 2 | 6 |
| *K. paraultunense* | MSLs |  |  |  | 47.8 ± 3 | 19.2 ± 3.2 | 27.2 ± 1.8 | 23.3 ± 3.8 | 25.6 ± 0.3 | 32 ± 2.5 |
|  | Lipids with ether bond |  |  |  | 12.8 ± 0.4 | 20.8 ± 1.7 | 24 ± 2.2 | 20.8 ± 2 | 21.4 ± 2.1 | 21.3 ± 1.7 |
| *S. acetigenes* | MSLs | 13.8 ± 3.7 | 26.8 ± 6.6 | 40.7 ± 9.3 |  |  |  | 16.6 ± 1.3 | 5.8 ± 2.6 | 6.9 ± 3.5 |
|  | Lipids with ether bond | 22.2 ± 2.4 | 21 ± 4.6 | 16.6 ± 3 |  |  |  | 9 ± 4.7 | 10 ± 3.2 | 7 ± 2.4 |

\* optimum growth temperature.
\*\* growth at different oxygen level was conducted at the optimal temperature of incubation for the two species

#### 2.3.1 Changes with growth temperature

We tested the potential changes in the cell biomass acid-hydrolysis lipid products of the two *Tissierales* strains at different growth temperatures. The relative abundance data of the acid-hydrolysed lipid products was determined as an average of *n*=3 biological replicates. When *K. paraultunuense* was grown at optimal temperature (55°C), 19 ± 3% of its hydrolysis-derived lipids were MSLs (Table 4; Supplementary Table 5 for full hydrolysis-derived lipid distribution). At 60°C this was 27 ± 2%, while at 45°C this was 48 ± 3%. The percent of lipids with an ether bond was stable (∼25%) at 55°C and 45°C and saw a slight increase to 32 ± 3% at 60°C (Table 4; Supplementary Table 5). In *S. acetigenes*, the hydrolysis-derived MSLs constituted 27 ± 7% of the total at optimum temperature (37°C; Table 4, Supplementary Table 6). At 47°C this increased to 41 ± 9%, while at 27°C this decreased to 14 ± 4%. The percent of lipids with an ether bond (23-26%) remained stable at all temperatures. These changes in both species suggest that at elevated temperatures, both species produce a higher relative abundance of MSLs. The opposite response of the two strains in their production of MSL temperatures below optimum suggests different membrane adaptative responses depending on the strain. The ether lipid high temperature response differed only slightly between the two strains. These lipid adaption differences may relate to the thermophilic character of *K. paraultunense* and the mesophilic character of *S. acetigenes* and may reflect evolutionary adaptations to distinct membrane composition strategies, or to strain specific thermal niches.

#### 2.3.2 Changes in the response to oxygen exposure

To assess the impact of oxygen exposure on the anaerobic cultures, cultures with a variable extent of oxygen exposure in the headspace were inoculated after three sub cultivation growth cycles and collected when cultures reached the exponential phase (between 0.08-0.1 OD_600_ for *S. acetigenes* and between 0.25-0.3 for *K. paraultunense*). When *K. paraultunense* was exposed to low amounts of oxygen in the headspace (1 and 2% O_2_), the percentage of hydrolysis-derived MSLs were 23 ± 4% and 26 ± 0.3% respectively. At 6% oxygen, this increased slightly to 32 ± 3% (Table 4; Supplementary Table 5), indicating a slight shift toward the production of MSLs components induced by the presence of oxygen. In *K. paraultunense* the percent of hydrolysis-derived lipids with an ether bond stayed very stable at all oxygen concentrations (21%) (Table 4; Supplementary Table 5).

In *S. acetigenes*, when oxygen was 1% the percentage of MSLs was 17 ± 1.3% (Table 4; Supplementary Table 6). At both 2 and 6% oxygen this decreased to 6 ± 3 and 7 ± 4%, respectively (Table 4; Supplementary Table 6). At all oxygen levels the percent of hydrolysis-derived lipids with an ether bond was again quite stable, 7-10% (Table 4; Supplementary Table 6). We conclude that the effect of oxygen on the production of the MSLs is again different depending on the strain of Bacillota tested, although they are both performing an anaerobic metabolism, for which the introduction of oxygen would be expected to create a similar effect.

The oxygen-dependent response in both species represent a specific system for membrane adaptation. While *K. paraultunense* increased the MSL production with increasing oxygen concentrations, *S. acetigenes,* exhibited a decrease in MSLs at all oxygen concentrations tested. On the other hand, oxygen exposure had almost no effect on the percent of lipids with an ether bond in both *K. paraultuense* and *S. acetigenes*. The strain-specific responses to environmental stressors, indicates that changes in bacterial community composition, could influence the distribution of MSLs in the environment. The effect of oxygen exposure on membrane spanning lipid production in members of the Acidobacteria has also been proven to be strain-related. Members of the Acidobacteria producing brGDGTs are either aerobic, microaerophilic or facultative anaerobic, thus known to be able to live in oxygen concentration lower than in the atmosphere (i.e., 21%). Acidobacterium *E. aggregans* was shown to stimulate brGDGT-Ia production at reduced oxygen concentration (i.e. 1% O_2_) (20) while the acidobacterium *Acidobacterium capsulatus* produced significant amounts of *iso*-DA after cell hydrolysis but lacked brGDGT (tetraether) at low O_2_ (20). On the other hand, the acidobacterium *Solibacter usitatus* Ellin6076 produced less brGDGTs per cell at 1% O_2_ (29). Both these results for Acidobacteria, and the ones obtained in the current study for anaerobic bacteria of the Bacillota phylum, support that the effect of oxygen exposure on brGDGT and brGDGT analogues is not the same and might involve different regulation mechanisms and adaptation processes. Overall, these results also give further support for a potential synthesis of brGDGTs and their building blocks by multiple bacterial types over a wide range of oxygen concentrations present in environmental systems.

### 2.4 Biosynthetic implications

We detected brGDGT analogues with two to four ester bonds, but not tetraethers (brGDGTs), in two members of the Bacillota phylum. Based on their chemical structures and their distribution (Fig. 4), we propose a theoretical biosynthetic scheme for brGDGT analogues in these two members of the Bacillota (Fig. 5). We hypothesize that these brGDGT analogues provide clues for our understanding of the bacterial biosynthesis of brGDGTs.

**Figure 5.**
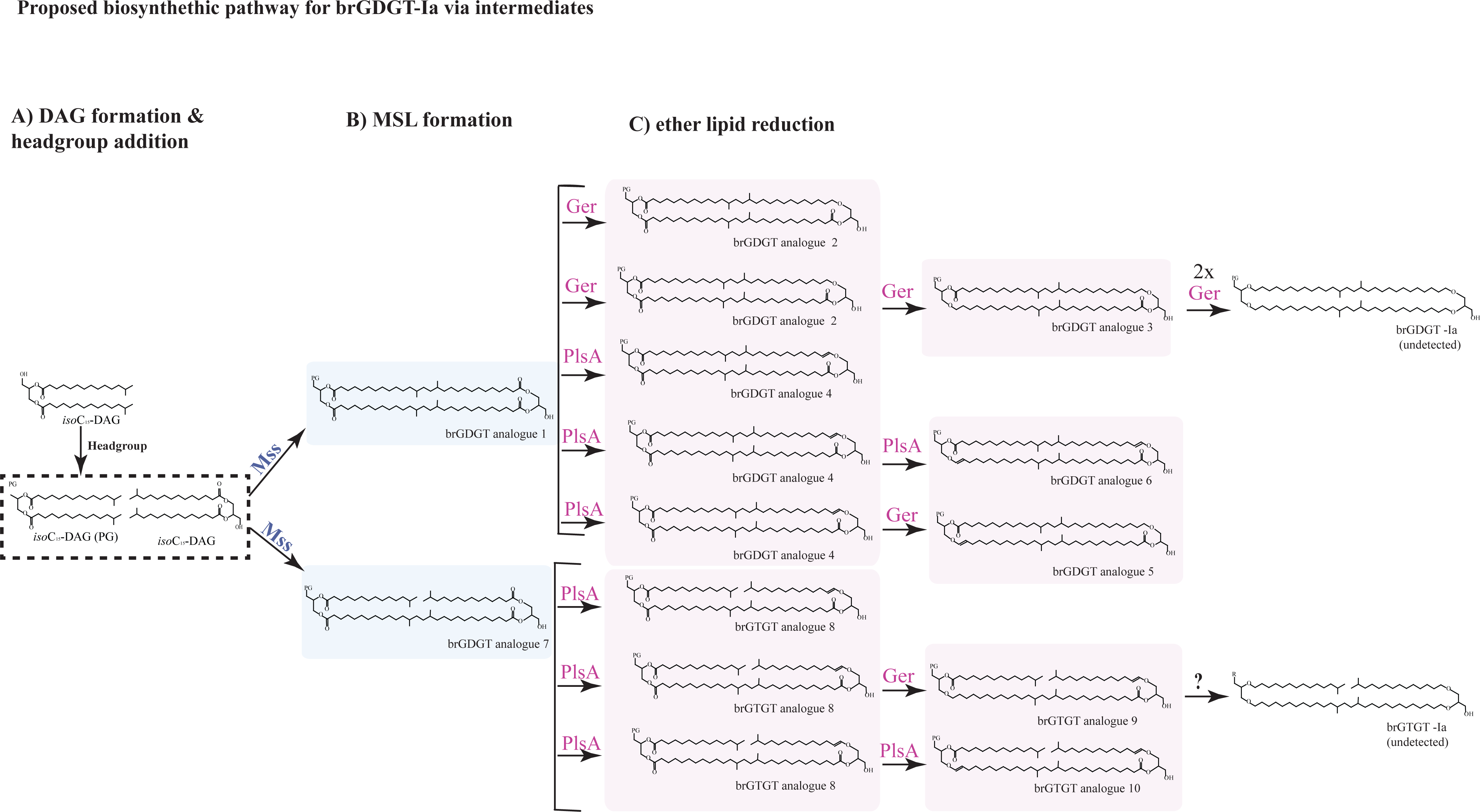
Proposed biosynthetic pathway for brGDGT and brGTGT analogues. Monomers of *iso-*C_15_-based DAG with or without a phosphatidylglycerol (PG) head group attached (A) would be condensed by the activity of Mss (B). The detection of brGTGT analogues suggests that the activity of Mss occurs before or after the formation of ether bonds (C), and that it could either be interrupted or proceed after the etherification process or happen simultaneously. We speculate that the etherification process (C) proceeds via the activity of PlsA and/or Ger (see main text for details). Branched GDGTs such as brGDGT-Ia or brGTGT-Ia have not been detected in the analyzed samples but we speculate here that could be formed by subsequent activity of Ger from brGDGT analogue 3, or the brGTGT analogues detected by the activity of a yet-unknown enzyme (indicated as a question mark). Alternatively, the etherification reactions could proceed on *iso-*C_15_-based DAG monomers with different levels of etherification; that would then be condensed via Mss (D). Here we have drawn all brGDGT analogues with an anti-parallel configuration, but it should be noted we have not determined whether this is the correct configuration. For all compounds, R = PG head group.

#### 2.4.1. Condensation reaction

In the current study, we have detected ester and ester/ether (vinyl and alkyl ether) bonded brGDGT and brGTGT analogues in two strains of Bacillota. Considering this finding and previous studies detecting ether/ester bonded membrane-spanning lipids both in *Thermotogota* (49, 54), and in other members of Bacillota (52, 55, 56), we conclude that the enzymatic reaction mediating the C-C condensation, catalyzed by Mss to form membrane spanning lipids, is not selective for the presence of ester or ether bonds when binding the lipid substrate (Fig. 5). This fact has been previously observed for the archaeal tetraether synthase Tes (57), enzyme that is able to bind ester-bonded phosphatidic acid in its active site, thus showing that Tes synthase can mediate its activity even in the presence of ester bonds linkages between the glycerol backbone and the fatty acid side chains (58) (Supplementary Figure 1).

#### 2.4.2. Head group involvement

The condensation to produce membrane-spanning compounds likely occurs after the head group has been added, as already suggested by our group (33). This hypothesis is further supported by the detection of the PG head group attached to one side of the brGDGT analogues in the Bacillota screened in this study, *Keratinibaculum* and *Sporoanaerobacter* (Fig. 3), as well as in the MSLs in *Thermotoga* (54), and in *Thermoanaerobacter* (31), which strongly suggests that diacylglycerol (DAG)-based MSLs biosynthesis proceeds from preformed DAG lipids, via a PG phospholipid intermediate (Fig. 5). In bacteria, PG is synthesized *de novo* starting from acylated glycerol-3-phosphate intermediates, containing a DAG core final assembly. Therefore, the attached PG moiety to the brGDGT analogue backbone, indicates that intact PG molecules serve as direct biosynthetic substrate in this pathway, where the head group from one side of the compound, is retained during the condensation process forming membrane-spanning lipids, being evidence of this the detection of *iso-*DA or alkyl and DMA C_30_ (Table 1, Supplementary Table 1).

Another line of evidence supporting the presence of a polar head group before the condensation reaction mediated by Mss is the similarity of its 3D structure with that of the archaeal Tes synthase. To this end, we conducted a superposition analysis of the bacterial Mss 3D models by AlphaFold (AF) and the AlphaFill (59) recovered ligands onto the Tes synthase (also known as GDGT-MAS) crystal structure, which showed that (i) the tested Mss enzymes conserve the same catalytic architecture as the Tes synthase, (ii) bacterial Mss can bind similar phospholipid ligands with polar head groups such as phosphatidic acid (LPP) and the archaeal lipid 2,3-di-*O*-phytanyl-*sn*-glycero-1-phosphate (L1P), as previously observed for Tes synthase (58), (iii) and the predicted presence of a SAM cofactor in the active site of both enzymes (Supplementary Fig. 1). This mechanistic interpretation will require future experimental confirmation.

#### 2.4.3. Synthesis of saturated alkyl ether or vinyl ether

Regarding the etherification process involved in the formation of the MSLs, the genomes of both the strains harbour several homologs of the *Pls*A-coding gene (Supplementary Table 7), coding for the PlsA enzyme involved in converting an ester bond in the *sn*-1 position of the glycerol backbone into a vinyl ether (plasmalogen) in a wide variety of bacteria and eukaryotes (60). The glycerol ester reductase Ger is a PlsA-like enzyme, converting ester to alkyl ether bonds at both the *sn*-1 and *sn*-2 positions of the glycerol backbone (33, 61). In general, a definite assignation of Ger or PlsA based only on the sequence primary structure is not possible. Although conserved sequence motifs have been described for the PlsA enzyme (60), the motif signature or the diagnostic functional features for the Ger enzyme remain to be determined and experimentally verified.

Several members of the Bacillota, previously screened in (33) and in the current study, synthesize plasmalogens (detected as DMAs) and in lower proportion saturated alkyl ethers, or only alkyl ethers, also harbor PlsA-like homologs with different functional domain architectures (33) (Supplementary Table 7). One of these homologs contains either one or two activation domains (*i.e.,* B, Pfam PF01869; Supplementary Table 7). For most of the genomes screened, the presence of only one activation domain B correlates with the production of plasmalogens and in a lower proportion of alkyl ethers, which we already speculated it could be due to the fact that one single activation domain B is potentially not enough to produce a fully functional Ger protein synthesizing alkyl ethers in higher proportion. In those cases, the synthesis of alkyl ethers could be a by-product in the synthesis of plasmalogens (33). We are basing this speculation on the fact that other microorganisms where only alkyl ether production had been detected, e.g. members of the Thermotogota and the Thermodesulfobacteriota (*Desulfatibaculum alkenivorans*), harbor two activation domains BB, in a PlsA-homolog which we annotate as an actual glycerol ester reductase Ger enzyme (33, 61), and actual validation of alkyl ether production through heterologous gene expression of those Ger coding sequences (33, 61).

In general, all the Bacillota genomes screened (Supplementary Table 7), contain another PlsA-homolog copy(s) but in those cases the activation domain is lacking, therefore it is unclear which role these enzymes might have. For the case of *K. paraultuense* and *S. acetigenes*, two PlsA-like homologs were detected in their genomes, which were in different operons (Supplementary Table 7). In both cases, one of them had two activation domains BB (2 x PF01869). We hypothesize that one of these homologs actually corresponds to the glycerol ester reductase Ger (33, 61), while the other homolog would be a PlsA catalyzing the formation of vinyl ethers (cf. Fig. 5C; Supplementary Table 7), working either independently or together (simultaneously or subsequently) in the reaction of ester reduction, pending of further confirmation.

In general, due to the low amount of ether bond-containing analogues, such as brGDGT analogue **2** and **3,** it is possible that the Ger enzyme activity is minor compared to that of the PlsA homologs in *K. paraultunense* and *S. acetigenes* under the growth conditions applied. Presumably, in other bacteria further sequential activity of the Ger enzyme to reduce the *sn*-2 position of the glycerol could ultimately lead to full reduction of ether bonds at the *sn*-2 positions and, therefore, lead to the synthesis brGDGT-Ia (not detected in *K. paraultunense,* nor in *S. acetigenes*). Our results indicate that the conversion of the ester bond to ether bond at *sn*-2 of the glycerol backbone is not performed or at least is not catalysed in *K. paraultunense* and *S. acetigenes* under the conditions tested here (i.e., IPL analysis was only carried out on biomass from cultures grown in optimal conditions). Future studies will need to address which conditions regulate the tuning system of the various PlsA/Ger homologs present in the genomes of these species responsible for the formation of the various types of ether bonds.

#### 2.4.4. Potential timing of biosynthetic steps

In our proposed scheme, Mss condenses PG-containing *iso-*C_15_-based DAG monomers (Fig. 5A) into branched glycerol diacyl tetraesters (brGDGT analogue **1**) (Fig. 5B). brGDGT analogue **1** can then be reduced by PlsA at the *sn-1* position to produce brGDGT analogue **4**, and subsequently brGDGT analogue **6** again by PlsA (Fig. 5C). Due to the high proportion of analogues **4** and **6** (cf. Fig. 4 and Table 3), it is possible that the PlsA enzymatic activity is increased in *K. paraultunense* and *S. acetigenes* or that these analogues are preferentially synthesized to adapt the membrane under the conditions of optimum growth in which the strains were grown. In a similar manner, Ger could reduce brGDGT analogue **1** at the *sn*-1 positions to produce brGDGT analogue **3 (**via brGDGT analogue **2)** (Fig. 5C). A similar pathway for etherification would then be expected for the synthesis of the brGTGT analogues, considering that brGTGT analogue **7** (Fig. 5B) **c**ould be transformed into brGDGT analogue **8** by the enzymatic activity of PlsA, and subsequently to brGTGT analogue **9** by Ger or alternatively to brGTGT analogue **10** by the enzymatic activity of PlsA again (Fig. 5C). As brGTGT analogue **10** is higher in relative abundance for the two strains (Fig. 4), this could suggest that this is the preferred product during optimal growth or that alternatively PlsA is more active in these strains at the tested growth conditions. We speculate that branched GDGTs, such as brGDGT-Ia or brGTGT-Ia, although not detected in the analyzed samples, may then be formed by subsequent activity of Ger from brGDGT analogue **3**, or in the case of the detected brGTGT analogues by the activity of a yet-unknown enzyme which could transform vinyl ethers into alkyl ethers, or by further etherification by Ger of alkyl ether-containing brGTGT analogues (not detected in this study) leading to brGTGT-Ia (Fig. 5C).

Our gathered evidence does not allow us to conclude as to whether the C-C condensation is mediated before the etherification process, after or simultaneously, although in Fig. 5B-C we have depicted the biosynthetic process with condensation by Mss prior to the etherification process. Alternatively, the etherification reactions by PlsA/Ger could proceed on *iso-*C_15_-based DAG monomers, that would then be condensed via Mss (Fig. 5D) leading to the different analogues mentioned above. Furthermore, the detection of brGTGTs (‘open GDGTs’) analogues with ether/ester bonds in *K. paraultense* and *S. acetigenes* suggests that the C-C condensation by Mss occurs either before or after the ester bonds are converted to ether bonds, and that it could either be interrupted or proceed after the formation of ether bonds (Fig. 5).

### 2.5 Implications for bacterial sources of brGDGTs in the environment

The large structural diversity and environmental distribution of brGDGTs suggests that these microbial lipids are produced by multiple bacterial groups (36). Recent studies have also suggested anaerobic heterotrophic bacteria as a source of brGDGTs in anoxic environments (36, 38, 41, 57). Our study has revealed that members of the phylum Bacillota, specifically of families including anaerobic heterotrophs, synthesize ester/ether brGDGT analogues, which could potentially be biotransformed into fully etherified tetraether brGDGT in the studied species under conditions not identified in this study or in other Bacillota species .

Recent papers described the occurrence of tetraesters (25, 48) and mixed ester/ether membrane-spanning lipids (48) with an *iso*-DA backbone containing additional methyl groups in mineral soils from Nepal and Rwanda (48), and in the Acidobacteria *Ca.* Solibacter usitatus (SD3) (25), which were hypothesized to be intermediate products of brGDGT biosynthesis. Acid hydrolysis of the polar fraction obtained from the extracts of the soils released *iso*-DA, *iso*-C_15_ and *iso*-C_17_ monoalkyl glycerol ethers (MGE) (48). As *iso*-C_15_ and *iso*-C_17_ MGEs had been previously found in members of the SD4 Acidobacteria (21, 22), the authors suggested Acidobacteria as potential biological sources of the detected ester/ether membrane spanning lipids based on *iso*-DA. Nevertheless, in our study we detected both the building blocks and ester/ether membrane spanning lipids brGDGT analogues, in members of the phylum Bacillota, confirming that bacterial groups other than Acidobacteria also synthesize these compounds, and potentially brGDGTs when exposed to different environmental conditions. Members of the phylum Bacillota are widespread in the environment, including soils, marine and freshwater systems, which is also compatible with their potential contribution to the pool of environmental brGDGTs.

### Conclusions

This study demonstrates, for the first time, that members of the Bacillota phylum produce branched GDGTs analogues, including ester/ether and vinyl ether-containing membrane-spanning lipids, thereby expanding the known diversity of potential bacterial sources of brGDGTs beyond Acidobacteria, and into the anaerobic bacteria realm. The production of these compounds, along with strain-specific responses to temperature and oxygen, suggests that brGDGT analogues synthesis is both environmentally regulated and taxonomically variable.

Together, our results indicate that brGDGT analogue biosynthesis in Bacillota proceeds via flexible, modular pathways in which C-C condensation by Mss can act on ester- and ether-linked phospholipid substrates, likely involving preformed phosphatidylglycerol intermediates and sequential conversion of ester groups into (vinyl) ethers by PlsA and Ger. These brGDGT analogues thus represent plausible biosynthetic intermediates that provide key mechanistic insight into how bacteria may synthesize canonical brGDGTs and tune membrane properties under specific growth conditions. These findings imply that brGDGTs may not be exclusive to Acidobacteria and reinforces the idea of the existence of biological sources of brGDGTs under different oxygen regimes.

## Material and Methods

### Genomic screening and search criteria

A screening of Mss and Ger proteins was performed in reference protein databases restricted to the Bacillota phylum. The protein sequences were searched against the BLAST database using the BLASTP algorithm and hits were considered as likely potential homologs based on a cut-off e-value ≤ 1e^-30^ and percentage of identity > 30%. For further selecting the Bacillota strains potentially producing brGDGT, a selection criterion was established; including that both the Mss and Ger proteins coding genes were both present in the genome, that the strain was available in pure culture, and that its isolation source was environmental.

### Strain cultivation

These strains were obtained from the German DSMZ culture collection and grown in their preferred media under anaerobic conditions. *Keratinibaculum paraultunense* was cultivated in 100% N_2_ at 55°C. The medium composed of (per liter) 5 g Trypticase peptone, 5 g meat peptone, 10 g yeast extract, 0.5 mL Sodium resazurin (0.1% w/v), 0.5 g L-cysteine HCl, 5 g Glucose, 40 mL salt solution. The salt solution was composed of 0.25 g CaCl_2_, 0.5 g MgSO_4_, 1 g K_2_HPO_4_, 1 g KH_2_PO_4_, 10 g NaHCO_3_, 2 g NaCl. *Sporanaerobacter acetigenes* was cultivated at 37°C in 80% N_2_ and 20% CO_2_. The medium composed of (per liter) 1 g NH_4_Cl, 0.3 g K_2_HPO_4_, 0.3 g KH_2_PO_4_, 0.2 g MgCl_2_, 0.1 g CaCl_2_, 0.1 g KCl, 0.6 g NaCl, 1 g yeast extract, 1.5 mL trace element solution SL-10, 0.5 ml Na-resazurin (0.1% w/v), 1.5 g Na_2_CO_3_, 2 g glucose, 0.5 g L-cysteine HCl, 0.3 g Na_2_S. Temperature experiments were carried out at 27, 37, 47 °C for *S. acetigenes* and 37, 45, 55 °C for *K. paraultunense*. Oxygen experiments were carried out by injecting a known volume (representing a 1%, 2% or 6% of the headspace) of sterile air into the completed medium. Oxygen was analysed using a PreSens Microx 4 trace with PSt7 Oxygen sensor. Before the experiments were inoculated, they were acclimatized twice in their respective condition. For *K. paraultunense*, cultures at 45°C were acclimatized twice before inoculated for the experiment. For the temperature 37°C, it was acclimatized once. The oxygen experiments were acclimatized for three transfers. Growth characterization was done in quadruplets whereas lipid analysis was done in triplicates. Growth was monitored by measuring their optical density at 600 nm (OD_600nm_). Cells were harvested at late exponential phase (OD_600nm_, 0.080-0.12 for *S. acetigenes* and 0.25-0.30 for *K. paraultunense*).

### Lipid extraction and analysis

Cells were collected from 60 mL of liquid culture by centrifugation, followed by freeze drying.

#### Hydrolysis-derived lipids

For hydrolysis-derived lipid analysis, the freeze-dried pellet was acid hydrolyzed with 2 mL of 1.4 N HCl in MeOH solution and refluxed for 3 h at 130 °C, after which the pH was adjusted to 4-5 with 2 N KOH in MeOH solution. Phase separation was done by the addition of 2 mL bidistilled water and 2 mL dichloromethane (DCM), and the organic layer was removed. The remainder was extracted two more times with DCM. The DCM layers were combined and dried after passing through Na_2_SO_4_ columns. The extracts were treated with a few drops of diazomethane prior to analysis to derivatize and methylate the acid groups. The alcohol groups were then trimethylsilylated using pyridine and N,O-bis(trimethylsilyl)trifluoroacetamide (BSTFA). Lipids were brought to a final concentration of 1 mg L^-1^ in ethyl acetate before analysis. Hydrolysis-derived lipid determination (relative abundance) was performed on an Agilent Technologies 7890B gas chromatography (GC) with a silica column (CP Sil-5; 25 m by 0.32 mm) and the samples were injected at 70°C. The oven temperature was programmed to 130 °C at 20 °C/min and then at 4 °C/min to 320°C, at which it was held for 10 min. The identification of hydrolysis-derived lipids was carried out on an Agilent Technologies 7890A GC coupled to an Agilent Technologies 5975C VL MSD mass spectrometer (MS) operated at 70 eV, with a mass range m/z 50 to 600 and 3 scans s^-1^. The column and oven settings were the same as those for the quantification GC analysis. Hydrolysis-derived lipids were identified based on literature data and library mass spectra. To aid with the identification of the long-chain DMAs, hydrolysis was repeated with deuterated methanol and analyzed by GC-MS as described above. Hydrolysis-derived lipids were also analyzed by reverse phase UHPLC-HRMS^n^ as described below.

#### Intact polar lipid extraction

Intact polar lipids (IPLs) were extracted from freeze-dried biomass using a modified Bligh and Dyer procedure and analyzed by reverse phase UHPLC-HRMS^n^ (62). An Agilent 1290 Infinity I UHPLC, equipped with thermostatted auto-injector and column oven, coupled to a Q Exactive Orbitrap mass spectrometer with an Ion Max source with a heated electrospray ionization probe (Thermo Fisher Scientific) was used. IPLs were separated on an Acquity BEH C_18_ column (Waters, 2.1 × 150 mm, 1.7 μm), maintained at 30°C, using a flow rate of 0.2 mL min^−1^. For this separation, a mixture of two different eluents was used; eluent A was composed of a mixture of methanol and H_2_O (85:15, v:v) and eluent B of methanol and isopropanol (50:50, v:v). Both eluents were modified by addition of small amounts of formic acid (0.12%, v/v) and 14.8 M NH3aq (0.04%, v/v). The elution program was as follows: 95% A for 3 min, followed by a linear gradient to 40% A at 12 min and then to 0% A at 50 min. These latter conditions were maintained until 80 min. The settings for the electrospray ionization probe, operated in positive ion mode, were: capillary temperature, 300°C; sheath gas (N_2_) pressure, 40 arbitrary units (AU); auxiliary gas (N_2_) pressure, 10 AU; spray voltage, 4.5 kV; probe heater temperature, 50°C; S-lens 70 V. The Q Exactive mass spectrometer was calibrated within a mass accuracy range of 1 ppm using the Thermo Scientific Pierce LTQ Velos ESI Positive Ion Calibration Solution. The IPLs were analyzed with a mass range of *m/z* 350–2000 with a resolving power of 70,000 ppm at *m/z* 200. Data-dependent tandem MS/MS (resolving power 17,500 ppm) was successively performed by fragmentation of the 10 most abundant ions (stepped normalized collision energy 15, 22.5, 30; isolation width, 1.0 *m/z*). Dynamic exclusion was used to temporarily exclude masses (for 6 s) to allow selection of less abundant ions for MS/MS. Note that IPLs have varying degrees of ionization efficiency. Hence, the peak areas (in response units) of different IPLs do not necessarily reflect their actual relative abundance. However, this method allows for comparison between samples when analyzed in the same batch.

---

**Supplementary Figure 1.**
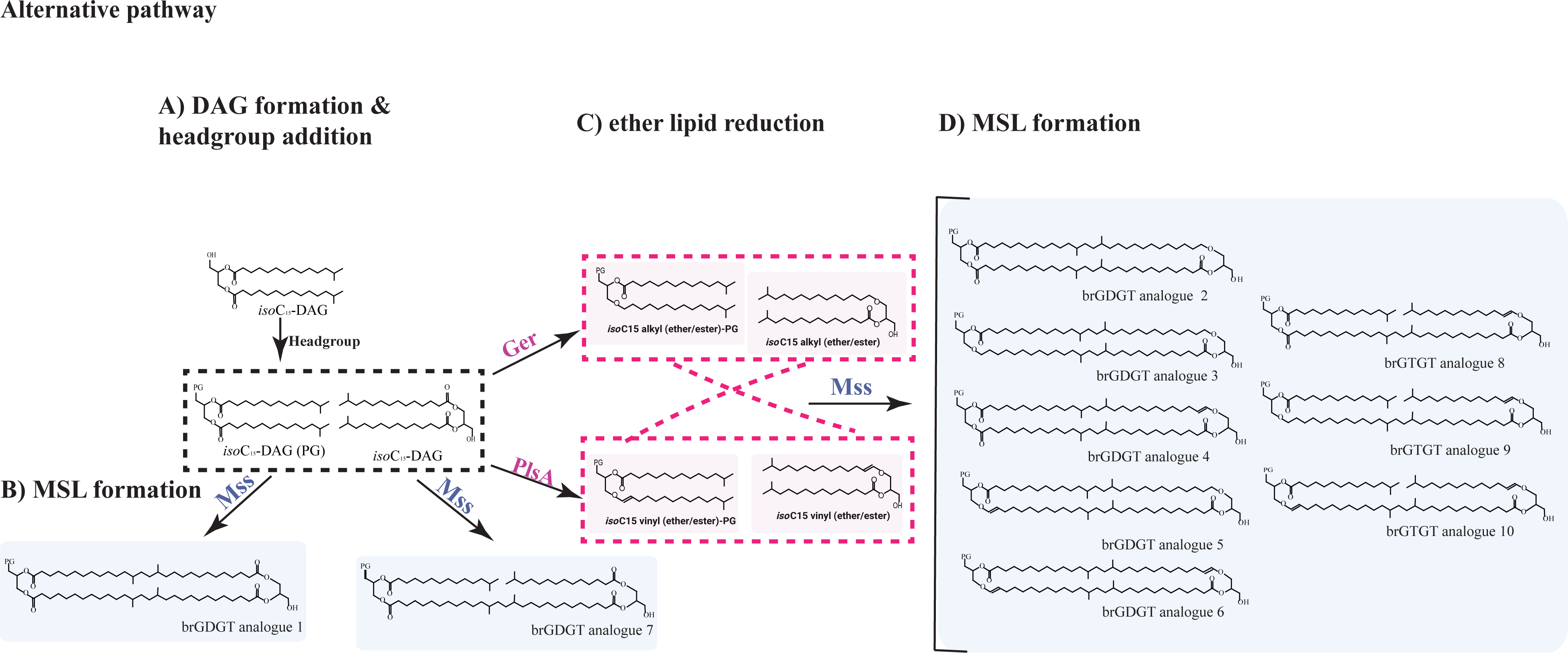
Architecture and tridimensional structure of the Membrane-spanning lipid enzymes from the archaeal Tes synthase (also known as GDGT-MAS; MJ0619) and bacterial Mss homologs. Comparison of the MJ0619 reference with bacterial Mss homologs. Left column: X-ray structure of Tes synthase/GDGT-MAS (58) shows the radical-SAM core containing three [4Fe–4S] (SF4) cofactors bound alongside lipid substrates LPP and L1P. Middle column: AlphaFold model of Mss from *K. paraultunense* (UniProt A0A4R3KZK4) incorporating AlphaFill SF4 cofactors and transplanted LPP/L1P substrates. Right column: AlphaFold model of Mss from *S. acetigenes* (UniProt A0A1M5Z5M3) with AlphaFill SF4 cofactors and LPP/L1P substrates. Proteins are displayed as cartoons; cofactors and ligands are shown as stick models. Bottom rows: Structural superposition onto GDGT-MAS (ChimeraX MatchMaker) demonstrates that both Mss enzymes position their radical-SAM [4Fe–4S] cluster and lipid substrates within similar hydrophobic tunnels, with the polar head groups extending toward the solvent interface. The conserved arrangement of cofactors and lipid tails indicates a common catalytic zone and substrate binding site consistent with the radical-SAM mediating tail-to-tail condensation of membrane lipid chains. LPP: Phosphatidic acid,L1P: 2,3-di-*O*-phytanyl-s*n*-glycero-1-phosphate.

## Acknowledgements

We thank Ilsa Posthuma for her technical support and help with anaerobic microbiology. We acknowledge Denise Dorhout, Jort Ossebaar and Monique Verweij for their technical support in lipid analysis. L.V. and J.S.S.D. received funding from the Soehngen Institute for anaerobic Microbiology (SIAM) and NIOZ.

